# The role of self-fertilization in plant colony establishment

**DOI:** 10.1101/2023.03.13.532419

**Authors:** Kuangyi Xu, Bryan Reatini

**Affiliations:** Department of Biology, University of North Carolina, CB#3280, Coker Hall, University of North Carolina, Chapel Hill, NC, USA; Department of Ecology & Evolutionary Biology, The University of Arizona, Tucson, AZ, USA

## Abstract

Increased rates of self-fertilization are often found in plant colonies, but the factors driving the observed higher selfing rates remain unclear. Specifically, the higher selfing rates in colonist populations may be due to 1) source populations with a higher selfing rate being more likely to successfully establish colonies (a filter effect), 2) the *in situ* evolution of selfing rate or a plastic selfing rate increase rescuing the colony from extinction, 3) selfing rate evolution post establishment. Using individual-based simulations and eco-evo models, we show that under both single and multiple dispersal, colony establishment may often be driven by a filter effect, due to a higher initial selfing rate and lower genetic load, which are correlated since selfing can purge deleterious mutations. Moreover, the role of the filter effect is weaker under multiple dispersal than single dispersal. The evolution of a higher selfing rate is unlikely to contribute directly to colony establishment. Although selfing rate evolution occurs during the colonization process, most of the selfing rate evolution may occur post establishment. Plasticity in selfing rates is more effective in facilitating colony establishment than the evolution of selfing.

## Introduction

The evolution of self-fertilization from predominantly outcrossing populations represents one of the most common transitions in flowering plant evolution (Barrett et al. 2008, Wright et al. 2008), and the mating system changes are often associated with environmental change (Eckert et al. 2010). Indeed, in colonizing plant populations, the breakdown of self-incompatibility (SI: the avoidance of selfed offspring through genetic recognition of shared alleles between parents) and an increase in selfing rates have been frequently reported (e.g., Reinartz and Les 1994, see Pannell 2015 for a review), while the maintenance of SI during colonization can lead to rapid population decline (e.g., DeMauro 1993, Colas et al. 1997). Moreover, biotas that are formed principally by colonization (e.g., oceanic islands) are enriched for self-compatible taxa (Grossenbacher et al. 2017).

Despite the evidence for increased rates of self-fertilization and the frequency of self-compatible taxa after colonization, the causal factor contributing to successful establishment and the enrichment of selfing on islands and island-like environments remains unclear (Cheptou 2019). There are several possible and mutually compatible explanations. The first explanation, as indicated by Baker (1955), is that colonization success should be more likely for individuals capable of self-fertilizing, as “a single propagule is sufficient to start a sexually reproducing colony”. This is often referred to as “Baker’s law” (Cheptou 2012, Pannell et al. 2015). Consequently, successful colonization by long-distance dispersal may filter for taxa with a sufficiently high selfing rate, which we refer to as the “filter effect”. A second explanation is that the selfing rate could evolve *in situ* during the process of colony establishment, and populations that fail to evolve a high selfing rate go extinct. In this case, successful colonization may depend on the rate of selfing evolution and thus the strength of selection on selfing. Alternatively, the evolution of a higher selfing rate may mainly occur post-establishment as a consequence of relaxation in the strength of ecological and genetic factors (e.g., inbreeding depression) that had inhibited the evolution of selfing during establishment. Finally, if the pollination conditions (e.g., pollen limitation) remain poor in the new habitat, a higher selfing rate on the island could result from mating system plasticity, that is, an immediate increase of the proportion of selfed seeds (Levin 2010).

Previous studies have identified several key genetic and ecological factors that affect the evolution of selfing. Selfing can evolve through a transmission advantage, where individuals with a higher selfing rate transmit more genes than those with a lower selfing rate since they can fertilize their own ovules in addition to those of other individuals. However, selfed offspring tend to have lower fitness relative to outcrossed ones, known as inbreeding depression (Charlesworth and Willis 2009). Classical models predict that a higher selfing rate is favored when inbreeding depression is lower than 0.5 (Llyod 1979, Lande and Schemske 1985, but see Goodwillie et al. 2005). Another important advantage of selfing is that selfing offers reproductive assurance when there is pollen limitation due to a lack of mates or pollinators (Ashman et al. 2004, Knight et al. 2005, Eckert et al. 2006). For this reason, selfing is expected to evolve in populations suffering from low population density or pollinator decline, such as those at the edge of a species range or in newly colonized environments (Levin 2012).

Mating system plasticity (hereafter referred as plasticity for brevity) has been widely documented in response to environmental stress (Steets and Ashman 2004, Travers et al. 2004, Ivey and Carr 2005, Good-Avila et al. 2008, Kay and Picklum 2013). Although an increased selfing rate through plasticity is quite intuitive, an important question is that to what extent can plasticity raise the probability of colony establishment. Previous individual-based simulations have shown that plasticity can greatly increase the probability of colony establishment (Peterson and Kay 2015), but the genomic mutation rate of deleterious mutations used in the simulation is small (∼0.01). Given that an instant increase of selfing rate can initially increase the genetic load due to exposure of deleterious mutations (Herlihy and Eckert 2002), the genomic mutation rate for deleterious mutations may profoundly influence the role of selfing on population survival. Therefore, it may be necessary to examine the role of plasticity in colonization by using a more realistic range of parameter values for deleterious mutation rates.

The lack of clarity on the role of selfing during colony establishment may be because the colonization process involves complicated genetic and demographic interactions. For example, the evolution of a higher selfing rate may increase the expression of deleterious mutations in homozygotes and result in a transient decrease in population fitness (Herlihy and Eckert 2002, Xu 2023), which presents a potential barrier to colonization success. For this reason, the fitness consequences of increased selfing during colonization may partly depend on the history of self-fertilization within the source population. Specifically, colonies dispersed from predominantly outcrossing source populations may be more susceptible to inbreeding depression than those formed from source populations since a history of self-fertilization will have purged some deleterious mutations. Additionally, since initial colonist populations are often small, the selfing rate, inbreeding depression, and genetic load of the colony will differ from those in the larger source population due to sampling effects (Kirkpatrick and Jarne 2000). Strong genetic drift can also have complicated effects on their later coevolution (Lynch et al. 1995, Bataillon and Kirkpatrick 2000). Moreover, since the strength of pollen limitation may depend on the population size, demographic changes will affect the intensity of selection on selfing, which in turn affects the strength of drift and the fitness, and thus the demographic growth.

Using individual-based simulations and eco-evo models, we investigate the roles of filtering effects, selfing rate evolution, and plasticity during colony establishment under one-time and multiple-time dispersal. We find that either a higher selfing rate of the source population or a higher initial selfing rate of the colony resulting from a bottleneck effect is crucial for colony establishment. Moreover, source populations with a higher selfing rate will purge more deleterious mutations and thus have a lower genetic load, which also contributes to colony establishment. For colonies initially under demographic decline, evolution of a higher selfing rate may facilitate extinction. For colonies that are initially growing, selfing rate evolution can only slightly increase the survival probability and is much less effective than mating system plasticity. Therefore, the evolution of selfing may more often be an unavoidable phenomenon accompanying the colony establishment process, rather than a driving force for establishment success.

## Methods

We adopt both individual-based simulations and an analytical eco-evo model with deterministic population genetics and stochastic demographic growth to interrogate how the selfing rate and its evolution contributes to colonization. The simulation is written in C++ and the code is available on Zenodo [submitted upon acceptance]. Key symbols used are summarized in Table 3.1. Instead of treating the selfing rate or fitness as a parameter, for both models, we allow the genetic load, inbreeding depression, and selfing rate to coevolve by incorporating both deleterious mutations and loci controlling the selfing rate (referred to as selfing modifiers hereafter).

**Table 1.**
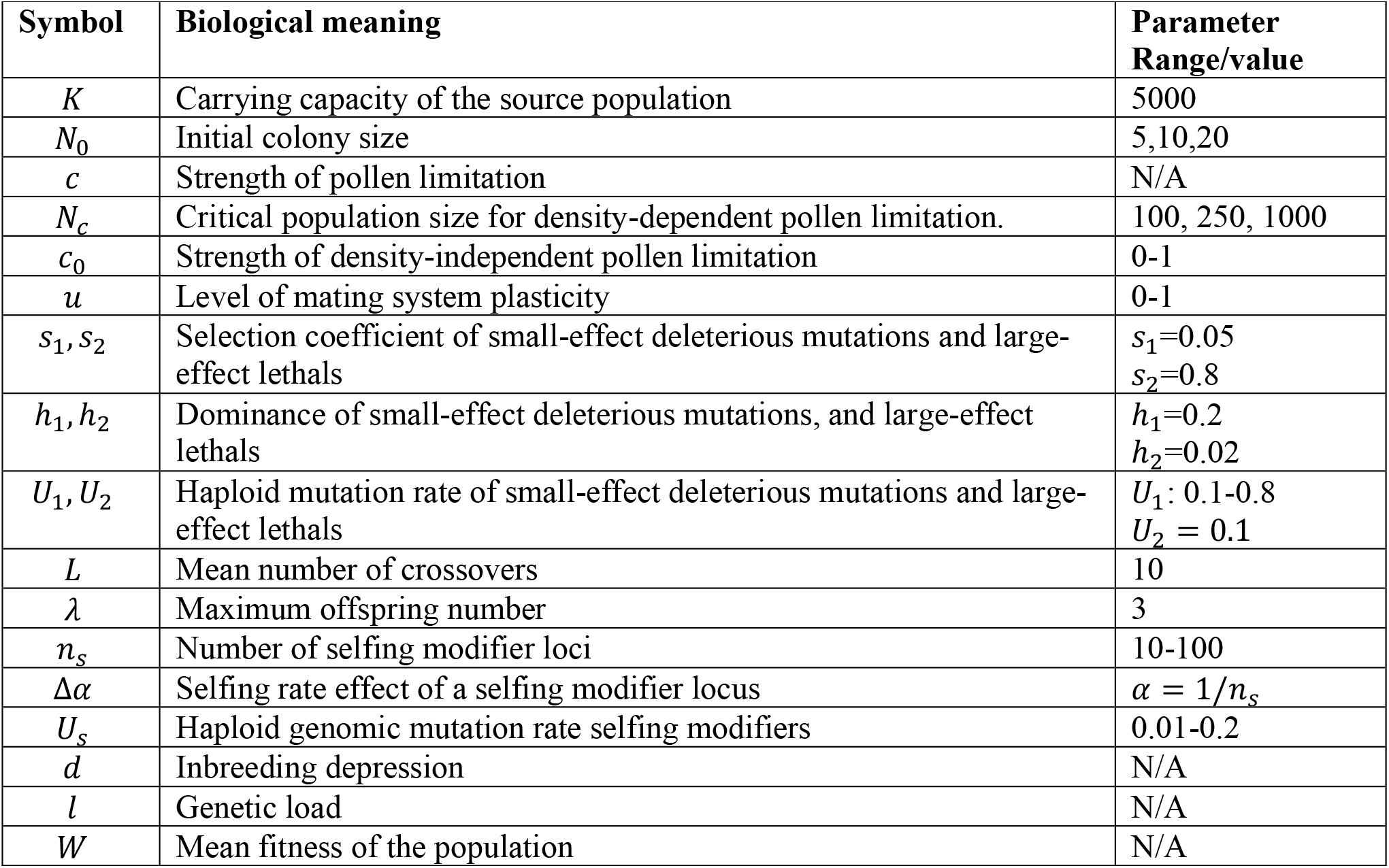
Biological meanings of key symbols used in the simulation and the model. The parameter range of some symbols is “N/A” because they are variables.

### Individual-based simulation

We model diploid, hermaphroditic populations with non-overlapping generations. The selfing rate of the population ranges from 0 to 1 based on the coevolution of selfing-modifiers and deleterious mutations described below, rather than being a binary-state trait (i.e., complete self-compatibility vs incompatibility). Each individual has two chromosomes with an expected number of crossovers *L* during meiosis. Deleterious alleles arise at an infinite number of loci along the chromosomes. Following empirical evidence (Charlesworth and Willis 2009, Eyre-Walker and Keightley 2007), we consider two categories of deleterious mutations. Mutations of the first category are small-effect, partially dominant deleterious mutations with selection coefficient *s*_1_ = 0.05, and dominance *h*_1_ = 0.2 (Mukai et al. 1972, Nakayama 2012). The second category consists of highly recessive lethal and semi-lethal mutations, with *s*_2_ = 0.8 and *h*_2_ = 0.02 (Simmons and Crow 1977, Klekowski and Godfrey 1989, Lande et al. 1994, Porcher and Lande 2005). The genomic mutation rates per chromosome of these two categories of mutations are *U*_1_ and *U*_2_, respectively.

We assume the selfing rate is determined by both genetic and plastic components, and the level of plasticity depends on the strength of pollen limitation in the environment. Considering the genetic component, empirically, the evolution of a higher selfing rate is often correlated with changes of a set of characters (e.g., anther-stigma distance), known as the “selfing syndrome” (van Kleunen and Ritland 2004, Ashman and Majetic 2006, Goodwillie et al. 2006, Fishman and Willis 2008, Goodwillie et al. 2010, Sicard and Lenhard 2011, Duncan and Rausher 2013). These mating-associated traits often have quantitative genetic basis with either few large-effect loci or many small-effect loci (Macnair and Cumbes 1989, Lin and Ritland 1997, Fishman et al. 2002, Goodwillie et al. 2006). These loci are thus effectively selfing modifiers as they regulate the selfing rate. We assume these selfing modifier loci have additive effects on the genetic value of the selfing rate. A number *n*_*s*_ of identical selfing modifier loci are evenly distributed along the chromosome, and each modifier locus changes the selfing rate effect by Δ*α*. Each modifier locus has two additive alleles *A* and *a*, and allele *A* is a selfing enhancer allele. We assume equal backward and forward mutation rates at each modifier locus as *μ*_*s*_ = *U*_*s*_/*n*_*s*_, where *U*_*s*_ is the haploid genomic mutation rate at selfing modifiers. Therefore, when there is no selection, the equilibrium selfing rate is 0.5. The equilibrium selfing rate is determined by the strength of inbreeding depression, pollen limitation and the net mutation pressure at selfing modifiers. An assumption of unequal mutation rates will only affect the critical strength of pollen limitation and inbreeding depression necessary for selfing to be selected for or against, but should not qualitatively change the role of selfing rate evolution during colony establishment.

Since the magnitude of the genomic mutation rate of selfing modifiers is unknown, in Appendix 3.2, we carry out a rough estimation of how large the genomic mutation rate of selfing modifiers can be. Briefly, we first use the unbiased estimator suggested by Zeng (1992) to estimate the effective number of loci (which tends to be a lower bound of the actual number of loci) underlying several mating-associated characters (e.g., anther-stigma distance, flower width), based on the data from the cross experiment of *Mimulus guttatus* and *M. cupriphilus* in Macnair and Cumbes (1989). We then calculate the genomic mutation rate of selfing modifiers by using the previous estimation of the base pair mutation rate in plants and the average number of base pairs per locus. However, it should be noted that this method assumes that every nucleotide substitution has a functional effect on the selfing rate, which can be an overestimation. We estimate the genomic mutation rate *U*_*s*_ to range from the order of 10^−4^ to 10^−1^ (see Supplementary Material), and this range is therefore used in our models.

For fecundity, each individual produces an identical number of ovules *λ*, which may be selfed, outcrossed, or unfertilized. Pollen limitation can be caused by density-dependent factors such as insufficient conspecific individuals (Hesse and Pannell 2011), or density-independent factors such as the lack of appropriate pollinators (Schueller 2004, Fishman and Willis 2008, Koski et al. 2017). We assume ovules used for selfing will always be self-fertilized, representing the advantage of reproductive assurance. It should be noted that in reality, the number of selfed ovules may also decline with increased strength of pollen limitation, since some modes of self-fertilization (e.g., geitonogamy, the self-fertilization caused by pollination between flowers of the same individual) require the facilitation of pollinators (Schoen and Lloyd 1992). Therefore, our model assumption maximizes the likelihood for selfing to facilitate colony establishment. We define the strength of pollen limitation *c* as a reduction in the probability for an ovule to be fertilized through outcrossing. The selfing rate of an individual when there is no plasticity is denoted by *α*. In this case, the number of selfed and outcrossed offspring is *αλ* and (1 − *α*)(1 − *c*)*λ*, respectively. The number of unfertilized ovules is (1 − *α*)*cλ*. To model plasticity, we assume individuals with a level of plasticity *u* will fertilize a proportion *u* of ovules remaining unfertilized, which thus produces an extra number of selfed offspring (1 − *α*)*cuλ* under pollen limitation strength *c*. This type of plasticity does not reduce the opportunity of outcrossing and thus always increases individual fitness, as may occur through delayed selfing (Goodwillie and Weber 2018). Note that there is another type of plasticity, not considered here, in which an increase in the number of selfed ovules comes at the cost of an opportunity for those ovules to outcross, such as through prior or competing selfing (Lloyd and Schoen 1992), but this form of plasticity is not considered in our simulations.

We model the strength of pollen limitation (Porcher and Lande 2005) as

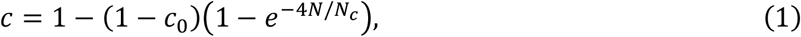

where *c*_0_ is the strength of pollen limitation caused by density-independent factors which is present even when the population size is large. The term 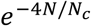 represents pollen limitation caused by density-dependent factors, where *N*_*c*_ is the critical population size that measures the sensitivity of pollen limitation to population size *N*. When *N* = *N*_*c*_, the effects of pollen limitation caused by population size is negligible.

We assume that populations with different genomic mutation rates of selfing modifiers *U*_*s*_ and deleterious mutations *U*_1_ have the same maximum offspring number, and *U*_*s*_ and *U*_1_ co-determine the selfing rates and fitness in the source population. For each combination of other parameters, we vary the values of *U*_1_ from 0.1 to 0.8 with an interval 0.1 (Mukai et al. 1972, Simmons and Crow 1977, Klekowski and Godfrey 1989), and the values of *U*_*s*_ from 0.01 to 0.2 with an interval 0.01. This gives a total of 160 parameter combinations of *U*_1_ and *U*_*s*_. For each parameter combination, the simulation occurs in two stages. In the first stage, a large source population evolves at the carrying capacity population size *K* for many generations to reach the mutation-selection-drift balance. After that, evolution of the source population stops, and colonization begins. The initial size of the colony is *N*_0_ and the migrants are randomly drawn from the source population without replacement. To assess the effects of selfing rate evolution on colony survival, we simulate two cases. In the first case, we fix the selfing rate of migrants to be a parameter, which is equal to the initial average selfing rate of the colony population. In the second case, we allow the selfing rate of migrants to evolve. During each replication, the simulation proceeds until either the colony goes extinct (i.e., *N* = 0), or is established (absolute fitness *≥* 1 and population size reaches the carrying capacity in the source population, *N ≥ K*). For both cases, we simulate 1000 replications and calculate the colony survival probability as the proportion of replicates for which establishment occurs.

Given that colonies formed by single-dispersal events most accurately represent those formed by long distance dispersal, we modify the simulations to allow for multiple dispersal to model proximate colonization events such as at the expanding edge of an invasion. For these simulations we allow new colonizing individuals to disperse to the colony every generation, these individuals are randomly drawn from the source population, and the number of migrant individuals each generation is drawn from a Poisson distribution with mean *N*_0_. Since under multiple dispersal, a colony will never become extinct, we limit the simulation for a maximum time of 500 generations. The survival probability is measured as the proportion of successfully established colonies during that time. We also record the time when each colony become established, and calculate the mean establishment time conditioned among established colonies. Due to this limited time frame, these simulations best represent rapid colonization events such as observed in biological invasions which occur during our limited window of observation as humans. Aside from these modifications, all other parameters remain the same between single and multiple dispersal simulations.

### Eco-evo model

Although the individual-based simulation is realistic in capturing both genetic and demographic stochasticity, this also makes it difficult to infer a causal relationship between the evolution of selfing and colony survival. Therefore, we build a simple deterministic eco-evo model to qualitatively investigate how the evolution of selfing affects the mean fitness and population size. The model only considers small-effect, partially dominant deleterious mutations. Large-effect lethal mutations contribute little to the genetic load since the genomic mutation rate is small (Mukai et al. 1972). The model captures the coevolution between the selfing rate, inbreeding depression and genetic load, but does not account for the accumulation of deleterious mutations. Therefore, the model tends to slightly overestimate fitness, since mutation accumulation is likely to occur when the population size becomes small (Charlesworth et al. 1993, Xu 2022a).

The effective selection coefficient of a selfing modifier allele with effect Δ*α* (Xu 2022b) is

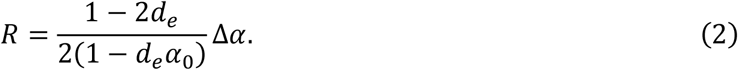

In equation (2), *d*_*e*_ is the effective inbreeding depression that incorporates both inbreeding depression *d* and pollen limitation *c*, given by

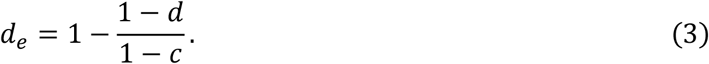

We denote the genetic load and inbreeding depression in the source population by *l*_0_ and *d*_0_, so that the absolute fitness of the source population is

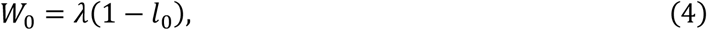

After colonization, the fitness of the colonizing population is

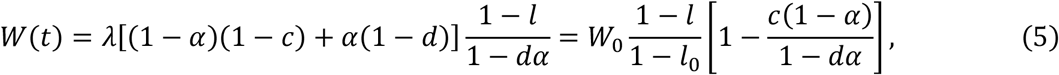

where the genetic load and inbreeding depression depend on the number of deleterious mutations *n*_1_ as in (Roze 2015)

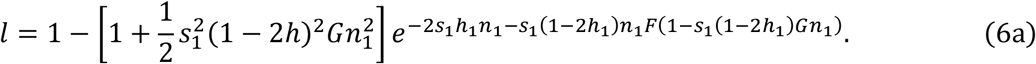

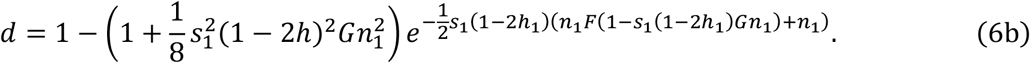

In equations (6a) and (6b), 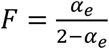 and 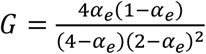 are the probability of identity by descent at one and two loci, respectively, where 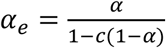 is the effective selfing rate after accounting for pollen limitation. The selfing rate depends on the frequency of selfing modifiers as

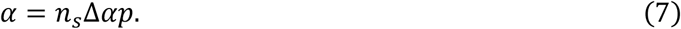

The fitness and selfing rate dynamics can be obtained by solving the recursion equations for the allele frequency of selfing modifiers *p*, and for the number of deleterious mutations per individual *n*_1_ as

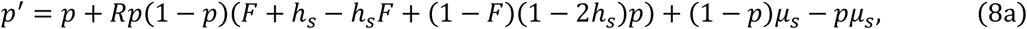

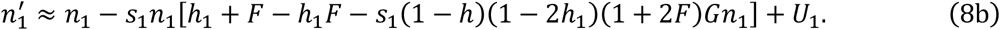

The selfing rate *α*_0_ and the number of deleterious mutations per individual *n*_0_ in the source population can be obtained based on equations (7), (8a) and (8b), by simultaneously solving 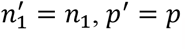.

After colonization happens, the initial absolute fitness of the colony population is

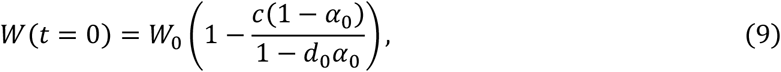

where *c* is given by equation (1) with *N* = *N*_0_. We treat the initial absolute fitness of the colony as a parameter value, and based on equation (9), we can solve the fitness of the source population *W*_0_. For the initial condition of the colony population, the expected initial frequency of the selfing modifier is equal to that in the source population 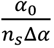, and the initial number of deleterious mutations per individual is *n*_1_(0) = *n*_0_. The population size changes as *N*(*t* + 1) = *W*(*t*)*N*(*t*), and the change of the absolute fitness *W*(*t*) can be obtained based on the recursion equations (8a) and (8b).

## Results

### Effects of the initial selfing rate and fitness on colony survival under single dispersal

In general, the selfing rate and absolute fitness of the source population strongly influences the colony survival probability. Figure 1(a) shows that there is a narrow range of the selfing rate and absolute fitness of the source population in which the colony survival probability is intermediate and rises quickly as the selfing rate or fitness increases. Below or above this critical range, the selfing rate and absolute fitness are either low or high, resulting in the colony survival probability to be either 0 or 1, respectively.

**Figure 1.**
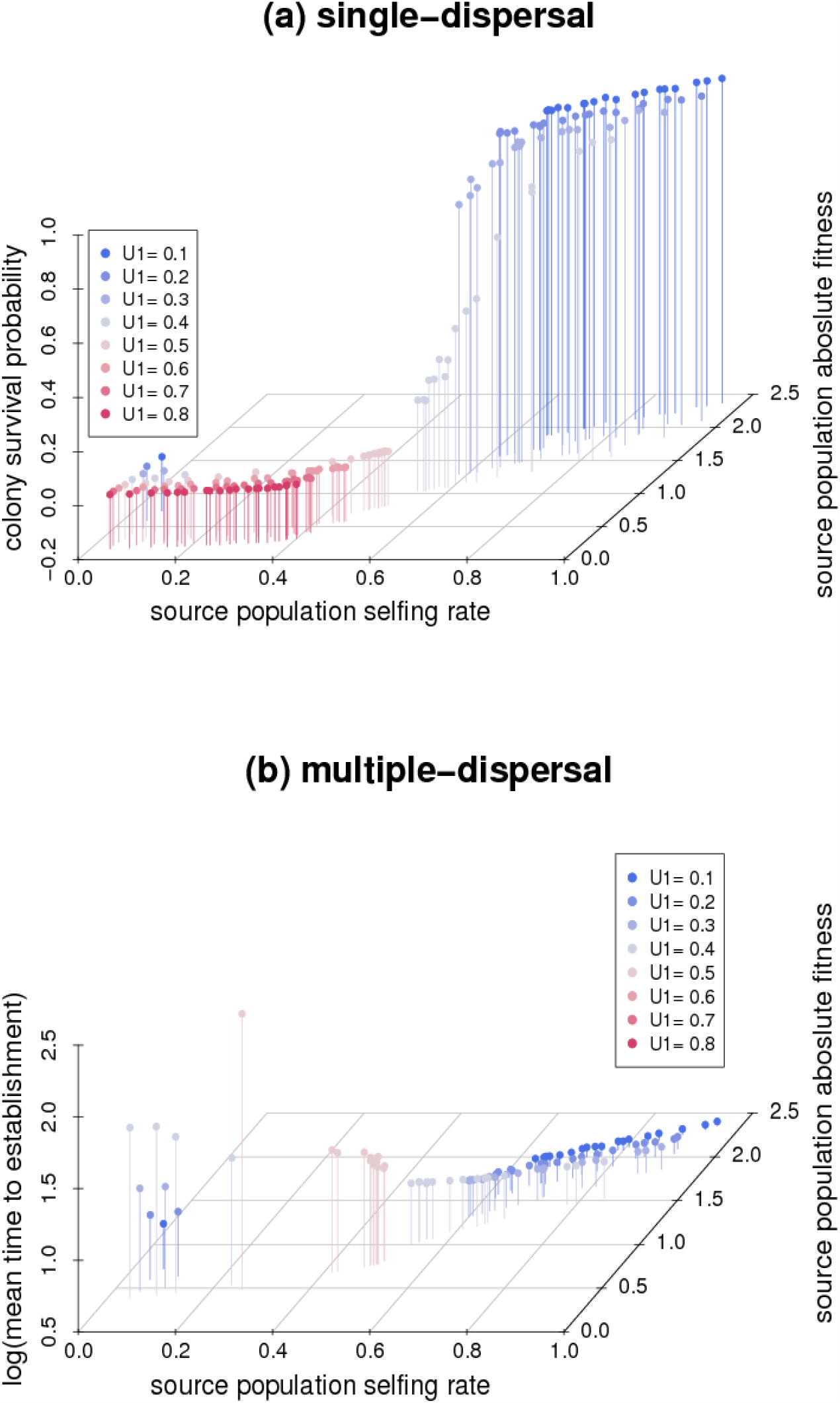
Effects of the selfing rate and absolute fitness in the source populations on the colony survival probability under single dispersal (panel (a)), and the mean time to establishment (in the log10 scale) under multiple dispersal (panel (b)). In both panels, dots with different colors have different *U*_1_, as indicated by the legend, and dots with the same color differ in the values of *U*_*s*_. Parameters are *K* = 5000, *L* = 10, *U*_2_ = 0.1, *n*_*s*_ = 10, Δ*α* = 0.1, *λ* = 3, *N*_0_ = 10, *N*_*c*_ = 100, *c*_0_ = 0, *u* = 0.

It should be noted that a higher selfing rate is positively correlated with a higher absolute fitness due to purging of deleterious mutations, as shown by the projection on the x-y surface in Figure 1(a). Populations with high genomic mutation rates of deleterious alleles *U*_1_ tend to have low selfing rates and low fitness (blue dots in Figure 1(a)), because a high *U*_1_ inhibits the evolution of selfing, and a low selfing rate in turn purges less deleterious mutations. On the other hand, populations with a low *U*_1_ tend to have high selfing rate and high absolute fitness, both contributing to colony survival.

### Eco-evo dynamics during colonization under single dispersal

Results from the eco-evo model indicate that an increase of the selfing rate cannot rescue colonies that are initially under demographic decline from extinction. Figures 2 illustrates the changes of the selfing rate, genetic load, and the population size relative to the initial colony size over time. Under density-dependent pollen limitation, the demographic decline rate is greater when the population size is smaller. As the black and grey solid lines in Figure 2(a) show, the decline in population size and fitness accelerates, since the initial demographic decline strengthens pollen limitation, which further reduces fitness in the short term. Moreover, as pollen limitation becomes strong, the selfing rate is effectively high due to there being almost no outcrossed ovules (even without selfing rate evolution), which results in a quick rise in the genetic load (see the dotted line in Figure 2(a) at *t* = 8). Strong pollen limitation also overcomes the strength of inbreeding depression and allows the selfing rate to evolve (see the dashed line at *t* = 9 in Figure 2(a)), but the selfing rate evolution exacerbates the rise in the genetic load (the uptick of the dotted line at *t* = 10 in Figure 2(a)). Although selfing rate evolution can purge the genetic load and increase the fitness later (after *t* = 10 in Figure 2(a)), there is no demographic opportunity for this to occur, since the population has already gone extinct by that time. Note that the results are still shown in this case because we adopt a deterministic genetic model. Under density-independent pollen limitation, as shown in Figure 2(b), a similar quick rise of the selfing rate and genetic load are still found (see *t*=60-70 in Figure 2(b)), although before that, the fitness decline is more linear (thus constant) compared to that under density-dependent pollen limitation.

**Figure 2.**
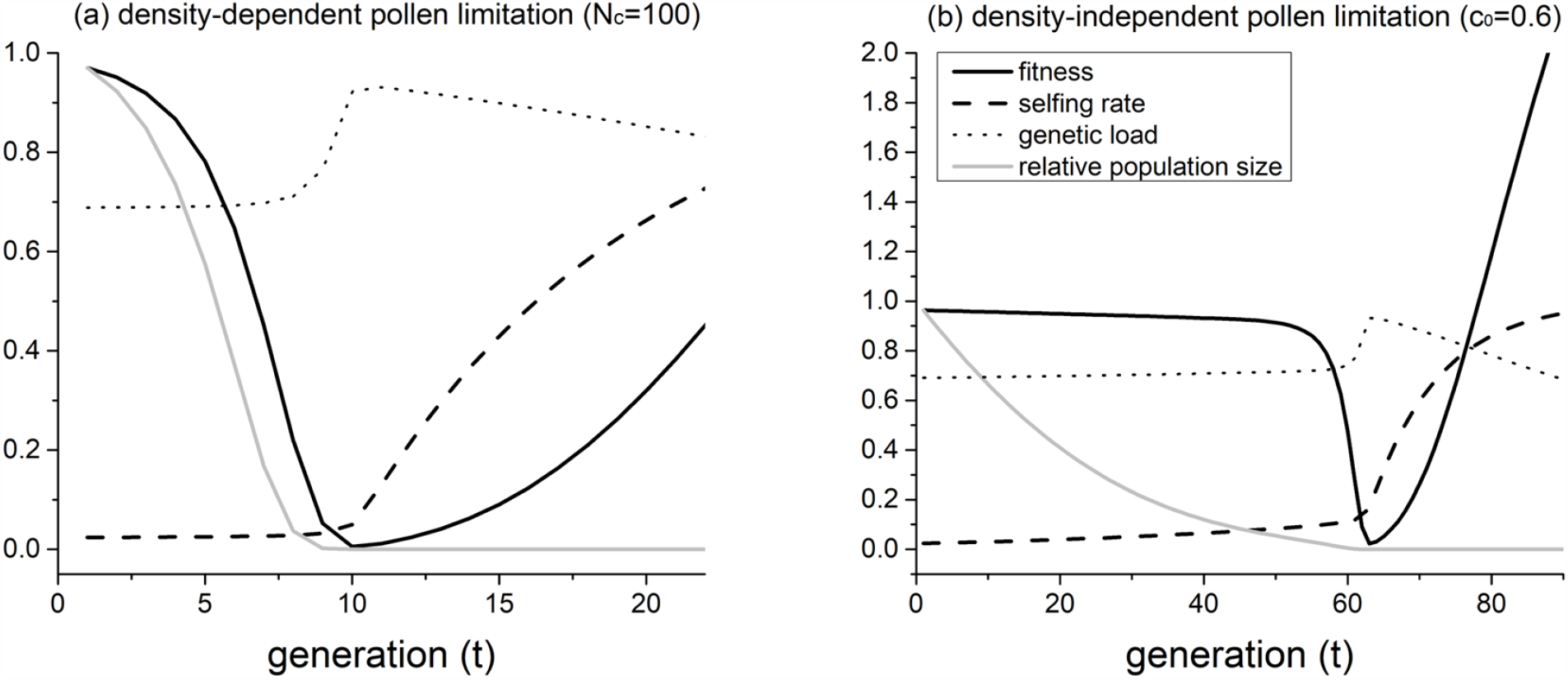
Temporal changes of the absolute fitness, selfing rate, genetic load, and the population size relative to the initial size *N*/*N*_0_. Parameters are *W*(0) = 0.98, *N*_0_ = 20, *n*_*s*_ = 10, Δ*α* = 0.1, *μ* = 10^−4^, *U*_1_ = 0.6, *u* = 0.

### The role of selfing rate evolution and plasticity on colony survival under single dispersal

The results from our eco-evo model indicate that selfing rate evolution is unlikely to rescue a colonizing population that is initially experiencing demographic decline because the rapid increase of the selfing rate will lead to a fitness-valley-crossing effect. As a result, we should expect that for a colonizing population to possibly survive, it should be initially growing. Indeed, in the individual-based simulations, for parameter combinations that give a non-zero colony survival probability, the initial colony growth rate is always positive (Figure S1). It should be emphasized that initially growing colonies still suffer a risk of extinction since the initial population size is small and there is demographic stochasticity. However, since our eco-evo model does not incorporate demographic stochasticity, it cannot tell us the role of selfing rate evolution on colony survival in initially growing colonies. Therefore, we turn to the results from individual-based simulation.

Analyses of individual-based simulation results suggest that the contribution of selfing rate evolution to colony survival is often very slight, unless pollen limitation occurs in a large population in the new habitat, which may be due to the lack of appropriate pollinators. Compared to colonies that have dispersed from the same population without selfing rate evolution, there are slightly more parameter combinations *U*_1_ and *U*_*s*_ that give non-zero survival probabilities for colonies with selfing rate evolution allowed (Table 2). Interestingly, when pollen limitation only occurs in small populations (*N*_*c*_ = 100), Figure 3 shows that the survival probability is negatively correlated with the rate of selfing rate evolution. Although one may suspect that the lower survival probability under quicker selfing rate evolution may be due to a temporary increase in the genetic load, it seems unlikely because the mean survival probability only differs slightly between colonies with and without selfing rate evolution (see the row “standard parameters” in Table 2). Therefore, the finding that colonies with lower survival probabilities have quicker selfing rate evolution may be a result of stronger selection on selfing due to more severe pollen limitation. However, such strong selection on selfing is still unable to rescue these declining populations.

**Table 2.**
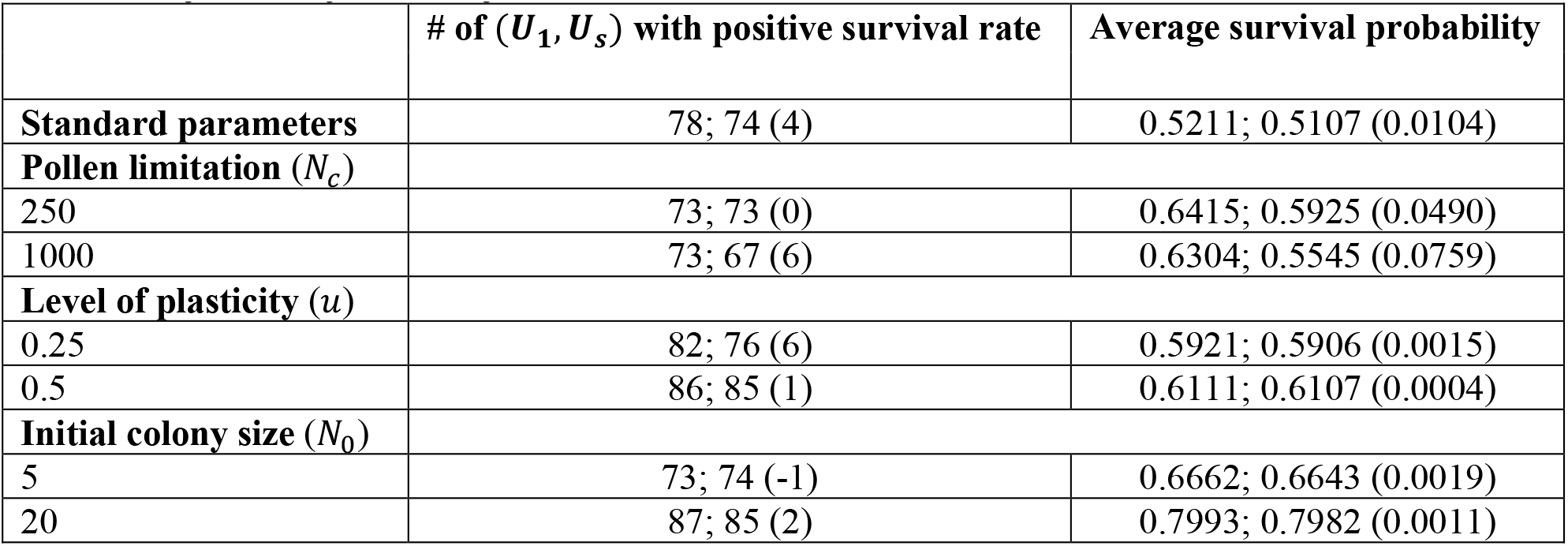
Comparison between colonies with the selfing rate allowed to evolve and fixed. The first column shows the number of parameter combinations (*U*_1_, *U*_*s*_) (out of a total of 160) that give positive colony survival rate. The second column shows the survival probability calculated by averaging over combinations (*U*_1_, *U*_*s*_) with colony survival rates between 0.05 and 0.95. In both columns, the first and second values are for colonies with evolving and fixed selfing rates, respectively, and the value in the parenthesis is their difference. The standard parameter values are *K* = 5000, *L* = 10, *U*_2_ = 0.1, *n*_*s*_ = 10, *λ* = 3, *N*_0_ = 10, *N*_*c*_ = 100, *c*_0_ = 0, *u* = 0.

| | # of ($U_1, U_s$) with positive survival rate | Average survival probability |
| --- | --- | --- |
| **Standard parameters** | 78; 74 (4) | 0.5211; 0.5107 (0.0104) |
| **Pollen limitation ($N_c$)** |  |  |
| 250 | 73; 73 (0) | 0.6415; 0.5925 (0.0490) |
| 1000 | 73; 67 (6) | 0.6304; 0.5545 (0.0759) |
| **Level of plasticity ($u$)** |  |  |
| 0.25 | 82; 76 (6) | 0.5921; 0.5906 (0.0015) |
| 0.5 | 86; 85 (1) | 0.6111; 0.6107 (0.0004) |
| **Initial colony size ($N_0$)** |  |  |
| 5 | 73; 74 (-1) | 0.6662; 0.6643 (0.0019) |
| 20 | 87; 85 (2) | 0.7993; 0.7982 (0.0011) |

**Table 3.** Comparison between survived and extinct colonies dispersed from the same population. The first and second column show differences between the survival and the extinction group. In the last two columns, the first and the second values are for the survival group and the extinction group, respectively. The standard parameter values are the same as those in Table 3.1. The values shown are the average across combinations of *U*_1_ and *U*_*s*_ that give the survival probability within the range 0.05-0.95.

|  | **Initial selfing rate difference** | **Initial genetic load difference** | **Rate of selfing rate evolution** | **Rate of genetic load change** |
| --- | --- | --- | --- | --- |
| **Standard parameter** | 0.0099 | -0.0332 | 0.0025; 0.0017 | 0.0036; 0.0266 |
| **Pollen limitation ($N_c$)** |  |  |  |  |
| 250 | 0.0146 | -0.0301 | 0.0045; 0.0004 | 0.0036; 0.0235 |
| 1000 | 0.0186 | -0.0259 | 0.0064; -0.0003 | 0.0041; 0.0218 |
| **Level of plasticity ($u$)** |  |  |  |  |
| 0.25 | 0.0057 | -0.3389 | 0.0015; 0.0006 | 0.0034; 0.0284 |
| 0.5 | -0.0003 | -0.0329 | 0.0008; -0.0001 | 0.0032; 0.0241 |
| **Initial colony size ($N_0$)** |  |  |  |  |
| 5 | 0.0196 | -0.0554 | 0.0022; -0.0022 | 0.0052; 0.0394 |
| 20 | 0.0015 | -0.0241 | 0.0014; 0.0027 | 0.0032; 0.0207 |

**Figure 3.**
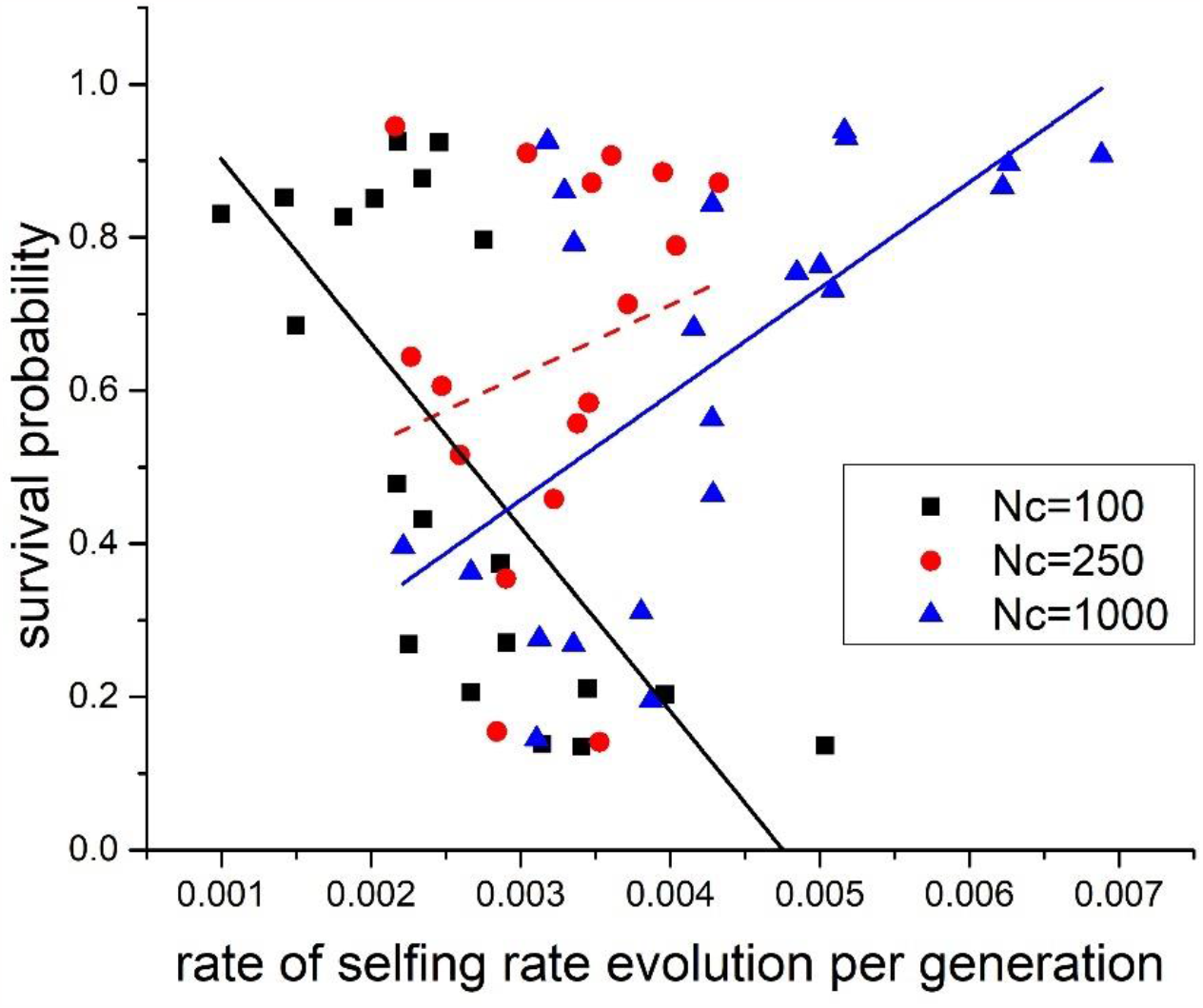
Relationship between colony survival probabilities and the rate of selfing rate evolution per generation. Each dot represents a combination of *U*_1_ and *U*_*s*_ that gives the colony survival probability between 0.05 and 0.95. Lines show results from linear regression, and solid lines showing significant relationships (p-value < 0.05), while the dashed line shows insignificant relationships. Other parameters are *K* = 5000, *L* = 10, *U*_2_ = 0.1, *n*_*s*_ = 10, Δ*α* = 0.1, *λ* = 3, *N*_0_ = 10, *c*_0_ = 0, *u* = 0.

In contrast, when pollen limitation is present even when the population size is relatively large (*N*_*c*_ = 1000), quicker selfing rate evolution is correlated with higher survival probabilities, as shown by blue lines and dots in Figure 3, and the mean survival probability differences between colonies with and without selfing rate evolution become prominent (see the row “*N*_*c*_ = 1000” in Table 2). Moreover, the contribution of selfing rate evolution to colony survival is small when the initial colony size is either small or large (compare the survival probability difference at *N*_0_ = 5,10,20 in Table 2). This is because drift is strong when the initial population size is too small, while when the initial colony size is relatively large, pollen limitation is weak under density-dependent pollen limitation.

Results from simulations that incorporate plasticity show that a higher level of plasticity *u* can promote colony survival. As shown by rows with *u* = 0,0.25,0.5 in Table 2, a larger *u* increases the number of parameter combinations *U*_1_ and *U*_*s*_ that give a non-zero survival probability. Moreover, plasticity can inhibit selfing rate evolution, which can be seen by comparing the rate of selfing rate evolution at rows with *u* = 0,0.25,0.5 in Table 3. This is because a higher background selfing rate reduces the selective strength on selfing modifiers (Xu 2022b).

### Comparison of colonies from the same population under single dispersal

For colonies dispersed from the same source population, how do successfully established colonies (the “survival group”) differ from colonies that go extinct (the “extinction group”)? By comparing these groups, we observed that even a slight initial difference in selfing rate and genetic load can lead to totally different colony fates. Specifically, the survival group generally has a higher initial selfing rate and lower initial genetic load than the extinction group (Table 3). Additionally, although for both extinction and survival groups, the genetic load builds up over time, compared to the extinction group, the survival group generally exhibits a higher rate of selfing rate evolution and a slower buildup of the genetic load (Table 3).

### Multiple dispersal

The previous subsections focus on the case of a single dispersal event, which is a reasonable model when the expected time interval between two colonization events is longer than the expected time for a colony to become established or extinct, as might be the case for a very remote island. When the colony is close to the source population, such as at the edge of an expanding range, multiple dispersal events could occur during the colony establishment process. Nevertheless, unlike the case of single dispersal, under multiple dispersal, the colony will either become established given enough long time, or never become established. Correspondingly, for nearly all (*U*_1_, *U*_*s*_) combinations, the colony establishment probability is either 0 or 1 (Figure S2). Since the colony survival probability is no longer a good measurement, we focus on the mean time to establishment in the log scale.

Generally, under multiple dispersal, the role of filtering, selfing rate evolution, and plasticity is qualitatively similar to what is found under one-time dispersal. As Figure 1(b) shows, a higher selfing rate and absolute fitness in the source population generally lead to quicker establishment, qualitatively consistent with the results under the single-dispersal case. Similarly, comparison of the mean time to establishment between colonies with selfing rates allowed to evolve versus fixed (Table S2) shows that evolution of selfing only slightly reduces the time to establishment, and its role is greater when strong pollen limitation can occur in large populations.

Interestingly, under multi-dispersal, populations with a low initial selfing rate and relatively low genomic mutation rate of deleterious mutations can successfully colonize given enough long time (blue dots with low selfing rate in Figure 1(b)). In contrast, under the case of single dispersal, colonies from the same population have low survival rate (blue dots with low selfing rates in Figure 1(a)). This may be because the demographic supplement in later generations under multiple dispersal can overcome the disadvantage of pollen limitation caused by a low selfing rate. Therefore, the initial selfing rate in the source population may be less important for colony establishment under multiple dispersal than under single dispersal.

## Discussion

Although increased rates of self-fertilization are widely found in colonizing populations, the underlying factors driving higher selfing rates observed in colonist populations remain unclear. This study addresses this question by investigating the role of selfing during colony establishment. For both scenarios of single and multiple dispersal, we show that successful colony establishment is mainly determined by a higher initial selfing rate and lower genetic load in the source population, which are correlated due to purging of deleterious mutations. Comparing colonies that have dispersed from the same source populations, we find that a slightly higher initial selfing rate and a lower initial genetic load are also crucial for colony survival. Both results suggest that a higher selfing rate in colonizing populations is mainly caused by a filter effect. This is consistent with what Baker’s Law suggests, although it should be noted that Baker’s original statement focuses on the compatibility of selfing instead of the rate of selfing (Baker 1955). Therefore, our results support the common assertion that the colonization process itself filters for populations with relatively high selfing rates.

We tested an alternative explanation for increased selfing rates in colonies, that evolution towards a higher selfing rate is crucial for colony establishment, but our results suggest that evolution of selfing only plays a slight role in facilitating colony survival. In fact, our eco-evo model suggests that for colonies initially experiencing demographic decline, the evolution of a higher selfing rate may facilitate extinction due to the exposure of deleterious mutations. Although an increase in the selfing rate can purge deleterious mutations over the long term, there is no demographic opportunity for this to occur when the population size is small, as is often the case for colonist populations. Nevertheless, our simulations show that the evolution of selfing can increase the colony survival probability (albeit slightly), particularly when the strength of pollen limitation is stronger.

Interestingly, for colonies dispersed from the same source populations, we find that compared to colonies that go extinct, colonies that survive have quicker evolution of the selfing rate and slower build-up of the genetic load. However, it is difficult to tell whether these are causes or consequences of colony survival. Specifically, colonies may survive because they happen to have a larger population size than those that go extinct due to demographic stochasticity. Since genetic drift is weaker in larger populations, larger populations will have quicker selfing rate evolution and slower accumulation of deleterious mutations, which may further contribute to a higher demographic growth rate and thus a larger population size. In this case, the evolution of selfing may simply be a phenomenon accompanying colony establishment, rather than a driving force for successful establishment itself. On the other hand, it may be also possible that the survival group may have quicker selfing rate evolution and a slower build-up of the genetic load due to the stochasticity of genetic drift, which results in a higher demographic growth rate.

If the evolution of selfing is a consequence of the colonization process, rather than its cause, then increased selfing rates observed in colonies may also in part be explained by the evolution of selfing after colony establishment. Although we do not run the colonist populations to equilibrium after establishment, the total selfing rate evolved during the colony establishment process is small (Figure S3), because the colony establishment or extinction process is fast. However, despite this small extent of selfing rate evolution during colonization, selfing evolution may result in a shift in the ultimate equilibrium state of selfing in the colony. Specifically, given that the genomic mutation rate of deleterious mutations is not too low, there exist two equilibrium states in our simulations, as classical models predict (e.g., Lande and Schemske 1985). One has a low selfing rate and high inbreeding depression, and the second one has a high selfing rate and low inbreeding depression. To which equilibrium the population will evolve depends on the initial selfing rate and inbreeding depression. Therefore, a small extent of selfing rate evolution and reduction in inbreeding depression due to strong genetic drift (Kirkpatrick and Jarne 2000) during colonization can cause the population to shift from the low-selfing-rate equilibrium to the high-selfing-rate equilibrium. Therefore, although our results suggest selfing rate evolution may not often contribute to colony establishment itself, it could still be an important contributor to the empirical observation of higher selfing rates in colonist populations.

Finally, for the role of mating system plasticity during colonization, we find that plasticity is more effective in promoting colony establishment than selfing rate evolution. However, plasticity can inhibit the effectiveness of a higher selfing rate in the source population and selfing rate evolution in promoting colony establishment, perhaps because plasticity reduces their importance in providing reproductive assurance. Moreover, plasticity, by increasing the background selfing rate, may reduce the selective strength on selfing modifiers and thus inhibit selfing rate evolution (Xu 2022b).

To facilitate ease of comparison and analysis, we made a number of simplifying assumptions in our models that should be addressed alongside their caveats. First, in order to compare populations with different parameters, we assume that all individuals produce the same number of ovules. In reality, populations with a higher selfing rate may produce more ovules, since selfing is often correlated with a smaller floral display and may save reproductive resources (Sato and Yahara 1999, Sicard et al. 2011). This extra fertility advantage of selfing populations thus can contribute to a higher establishment ability. Nevertheless, higher selfing rates are correlated with lower dispersal ability (Cheptou and Massol 2009, Massol and Cheptou 2011). As a result, although populations with a higher selfing rate may have a higher colony establishment ability, colonies may not necessarily have an overrepresentation of selfing populations. However, it should be noted that the evidence that dioecious species are enriched on islands (Carlquist 1966, Baker and Cox 1984, Webbet al. 1999, Schlessman et al. 2014) does not necessarily support that outcrossers are more likely to be successful long-distance colonizers, since the evolution of dioecy may be a result of selection to avoid selfing after colony establishment (Grossenbacher et al. 2017).

Despite the above assumptions, our models allow us to draw a few conclusions regarding the causal factors driving higher selfing rates observed in colonist populations. First, our results show that successful colony establishment may often be driven by a filtering effect on source populations, consistent with Baker’s Law. Second, plasticity in selfing may broaden the range of source populations that may pass this filter and may play a more important role than the evolution of selfing in facilitating colony establishment. Finally, although the evolution of selfing rate is unlikely to contribute directly to colony establishment, it does indeed occur during the colonization process and may therefore contribute to an enrichment of selfing within colonist populations if it results in a shift in equilibrium states.

## Notes

### Competing Interest Statement

The authors have declared no competing interest.

