## Supplementary Material for "The role of self-fertilization in plant colony establishment"

**Supplementary Methods**

Here we present a rough estimation of how large the genomic mutation rate of selfing modifiers can be in wild populations. We first estimate the number of genes underlying several mating-associated characters, using the data from the cross experiment of *Mimulus guttatus* and *M. cupriphilus* in Macnair and Cumbes (1989). We use the unbiased estimator suggested by Zeng (1992) to estimate the actual number of loci for each floral character as

$$n = \frac{2\bar{c}n_e + C_\alpha(n_e - 1)}{1 - n_e(1 - 2\bar{c})} \qquad \text{(S1)}$$

$n_e$ is the effective number of loci from Castle-Wright estimation (Lynch and Walsh 1998, Chapter 9), which assumes all loci are identical. The parameter $\bar{c}$ measures recombination between loci and is estimated by $\bar{c} = (M - 1)/2M$, as suggested in Lynch and Walsh (1998), where $M$ is the haploid chromosome number. Since $M = 14$ for the two *Mimulus* species, we get an estimation of $\bar{c} = 0.47$. $C_\alpha$ is the squared coefficient of variation of additive effects and is hard to estimate. However, analysis of data from mutation accumulation experiments of *Drosophila melanogaster* in Keightley (1994) shows that $C_\alpha$ is on the order of 6, 17, and 24 for several characters. These high values of $C_\alpha$ suggests a leptokurtic distribution of allelic effects, with a high density of small-effect alleles. Therefore, to estimate the range of number of loci, we calculate the actual number of genes $n$ with different $C_\alpha$ values at $0.5$,10, and 24 which are chosen to give the lower and upper limit.

Table S1 summarizes the estimated numbers of loci for the 11 characters measured in Macnair and Cumbes (1989) based on equation (S1). Characters that are most related to selfing rate, like stamen length, style length and stigma-anther distance, are usually controlled by a larger number of genes. By assuming no pleiotropy, across the 11 mating-associated characters, the total number of genes is on the order ranging from $3 \times 10^2$ to $5 \times 10^3$. Since previous research has found the base pair mutation rate in plants to be on the order of $3 \times 10^{-10}$ to $7 \times 10^{-9}$ (Ossowski et al. 2010, Ness et al. 2012, Keightley et al. 2015), assuming 1000 bp per locus (Juenger et al. 2000), the lower bound of the genomic mutation rate is $(3 \times 10^2) \times 1000 \times (3 \times 10^{-10}) \sim 10^{-4}$, and the upper bound is $(5 \times 10^3) \times 1000 \times (7 \times 10^{-9}) \sim 10^{-1}$. Therefore, the haploid genomic mutation rate $U_s$ across the 11 characters is estimated to be on the order of $10^{-4}$ to $10^{-1}$. This may be an under-estimate, as previous direct estimates of the genomic mutation rate of quantitative characters often give higher values. For example, the haploid mutation rate per character in *Drosophila melanogaster* for bristle traits is estimated to be about 0.015 (Mackay et al. 2005). Estimation of the haploid genomic mutation rate for the square root of total fruit production in a wild population of *Arabidopsis thaliana* gives a high value in the range of about 0.08-0.13 or 0.15-1.0 based on two different parameter-estimation methods (Rutter et al. 2010).

### Supplementary Tables and Figures

**Table S1:** Estimation of numbers of genes underlying some mating-associated traits using the data from Macnair and Cumbes (1989). $n_e$ is the effective number of loci predicted from the Castle-Wright estimator, assuming all the loci are identical. $n$ is the actual number of loci predicted by the unbiased estimation equation (S1) under different values of $C_\alpha$.

| **Character** | $\boldsymbol{n_e}$ | $\boldsymbol{n(C_\alpha = 0.5)}$ | $\boldsymbol{n(C_\alpha = 10)}$ | $\boldsymbol{n(C_\alpha = 24)}$ |
|---|---|---|---|---|
| Flower width | 6 | 14 | 95 | 214 |
| Flower height | 4 | 6 | 38 | 85 |
| Pistil length | 11 | 49 | 353 | 800 |
| Ovary length | 3 | 4 | 23 | 51 |
| Style length | 12 | 65 | 469 | 1064 |
| Corolla length | 6 | 14 | 95 | 215 |
| Lower corolla lobe width | 4 | 7 | 45 | 100 |
| Lower corolla lobe length | 5 | 9 | 62 | 140 |
| Mean stamen length | 10 | 35 | 248 | 561 |
| Stigma-anther distance | 14 | 125 | 913 | 2073 |
| Number of spots on corolla | 3 | 5 | 31 | 69 |
| **Total** | 79 | 333 | 2370 | 5373 |

**Table S2.** Comparison between colonies with the selfing rate allowed to evolve vs. fixed in the scenario of multiple dispersal. The columns show the mean value of $\log_{10}$(colony establishment time) averaged over parameter combinations $(U_1, U_s)$ (out of a total of 160) with establishment probabilities greater than 0.05. The standard parameter values are $K = 5000, L = 10, U_2 = 0.1, n_s = 10, \lambda = 3, N_0 = 10, N_c = 100, c_0 = 0, u = 0$.

| | **Mean $\log_{10}$(establishment time) (selfing rate evolves)** | **Mean $\log_{10}$(establishment time) (selfing rate fixed)** | **difference** |
|---|---|---|---|
| **Standard parameters** | 0.8345 | 0.8354 | -0.0010 |
| **Pollen limitation** ($N_c$) | | | |
| 250 | 0.9275 | 0.9235 | -0.0040 |
| 1000 | 0.8499 | 0.8923 | -0.0424 |
| **Level of plasticity** ($u$) | | | |
| 0.25 | 0.8216 | 0.8240 | -0.0024 |
| 0.5 | 0.8114 | 0.8145 | -0.0031 |

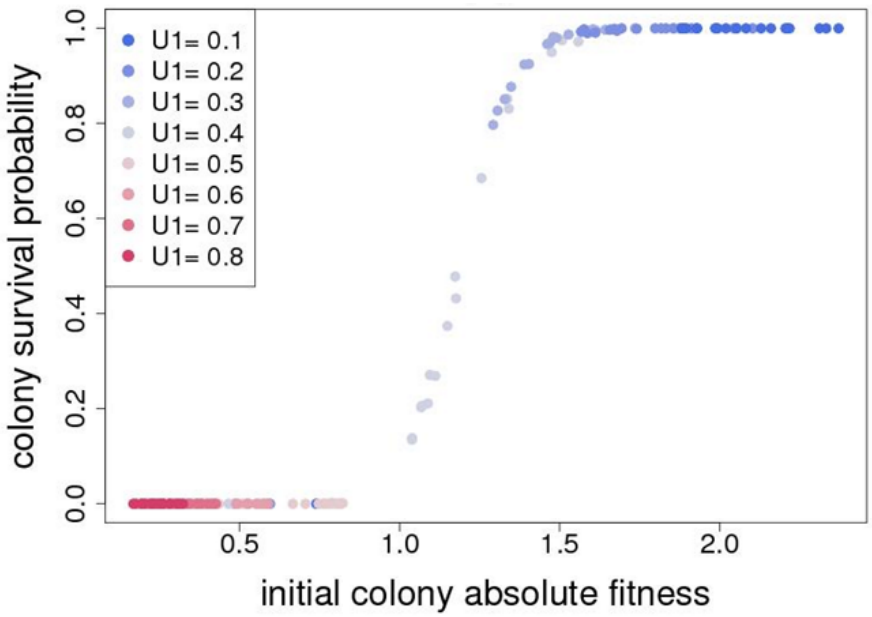


**Figure S1.** Effects of the mean initial absolute fitness of colonies dispersed from the same source populations on the colony survival probability. Legends and parameters are the same as those in Figure 1.

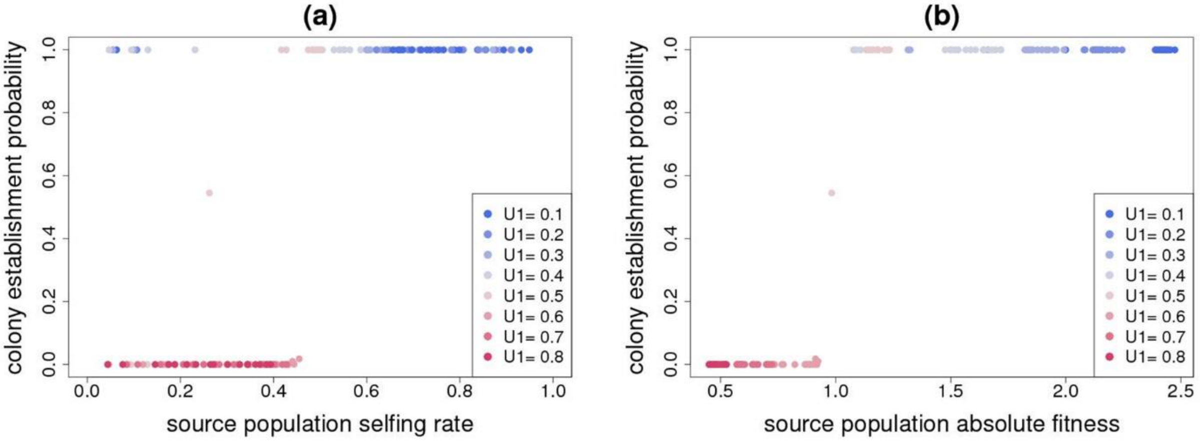


**Figure S2.** The probability of colony establishment before 500 generations in the scenario of multiple dispersal. Parameters are the same as those in Figure 1.

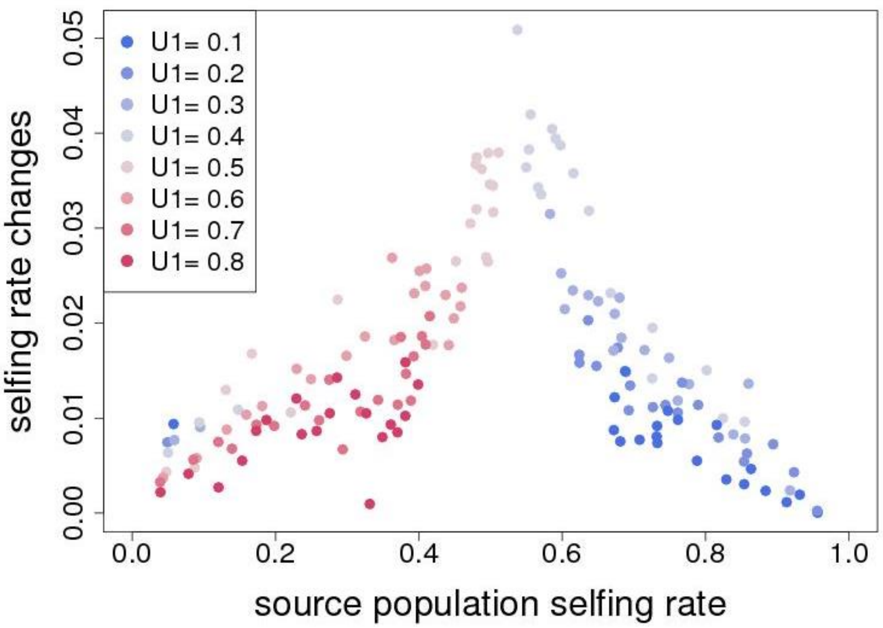


**Figure S3.** The overall evolutionary changes of the selfing rate of colonies upon colony extinction or establishment. Legends and parameters are the same as those in Figure 1.
